# Prolonged XPO1 inhibition is essential for optimal anti-leukemic activity in *NPM1*-mutated AML

**DOI:** 10.1101/2021.12.11.472216

**Authors:** Giulia Pianigiani, Andrea Gagliardi, Federica Mezzasoma, Francesca Rocchio, Valentina Tini, Barbara Bigerna, Paolo Sportoletti, Simona Caruso, Andrea Marra, Giulio Spinozzi, Sharon Shacham, Yosef Landesman, Concetta Quintarelli, Franco Locatelli, Maria Paola Martelli, Brunangelo Falini, Lorenzo Brunetti

**Affiliations:** Hematology, Department of Medicine and Surgery, Center for Hemato-Oncological Research (CREO), University of Perugia, Perugia, Italy; Institute of Hematology and Bone Marrow Transplantation, Santa Maria della Misericordia Hospital, Perugia, Italy; Department of Hematology and Oncology, Cell and Gene Therapy, IRCCS Bambino Gesù Children’s Hospital, Rome, Italy; Karyopharm Therapeutics, Newton, MA, USA; Department of Translational Medical Sciences, Federico II University, Naples, Italy; Department of Gynecology/Obstetrics & Pediatrics, Sapienza University of Rome, Rome, Italy

## Abstract

*NPM1* encodes for a nucleolar multifunctional protein and is the most frequently mutated gene in adult acute myeloid leukemia (AML). *NPM1* mutations cause the aberrant accumulation of mutant NPM1 (NPM1c) in the cytoplasm of leukemic cells, that is mediated by the nuclear exporter Exportin-1 (XPO1). Recent work has demonstrated that the interaction between NPM1c and XPO1 promotes high homeobox (HOX) genes expression, which is critical for maintaining the leukemic state of *NPM1*-mutated cells. However, the XPO1 inhibitor Selinexor administered once or twice/week in early-phase clinical trials did not translate into clinical benefit for *NPM1*-mutated AML patients. Here, we demonstrate that this dosing strategy results in only temporary disruption of the XPO1-NPM1c interaction and transient HOX genes downregulation, limiting the efficacy of Selinexor in the context of *NPM1*-mutated AML. Since second-generation XPO1 inhibitors can be administered more frequently, we compared intermittent (twice/week) versus prolonged (5 days/week) XPO1 inhibition in *NPM1*-mutated AML models. Integrating *in vitro* and *in vivo* data, we show that only prolonged XPO1 inhibition results in stable HOX downregulation, cell differentiation and remarkable anti-leukemic activity. This study lays the groundwork for the accurate design of clinical trials with second-generation XPO1 inhibitors in *NPM1*-mutated AML.

## Introduction

*NPM1*-mutated acute myeloid leukemia (AML) accounts for about one third of AML in adults^1,2^. The most distinguishing feature of *NPM1*-mutated cells is the aberrant localization of mutant NPM1 (NPM1c) in the cytoplasm^1^, caused by the loss of a nucleolar localization signal and the gain of a nuclear export signal within the C-terminal end of NPM1^3,4^. Both NPM1c nuclear export and cytoplasmic accumulation are dependent on its interaction with the nuclear exporter Exportin-1 (XPO1)^3,4^. Another unique property of *NPM1*-mutated AML is the high expression of homeobox (HOX) genes and their cofactors *MEIS1* and *PBX3* (hereafter referred as to HOX/MEIS)^5,6^. We recently found that high HOX/MEIS levels are required to maintain the undifferentiated state of leukemic cells^7^ and that HOX/MEIS expression is directly dependent on the interaction between NPM1c and XPO1^7^.

The selective inhibitors of nuclear export Selinexor and Eltanexor covalently bind XPO1 and disrupt the interaction with its cargo proteins^8^, including NPM1c^7^. Preclinical studies have demonstrated that XPO1 inhibition cause NPM1c nuclear relocation, loss of HOX expression, differentiation, and growth arrest of *NPM1*-mutated cells^7,9,10^. However, patients with *NPM1*-mutated AML showed suboptimal responses to Selinexor in early-phase clinical trials^11–14^. As Selinexor has a half-life of 6 hours^11^ and was administered once or twice/week, we hypothesized that intermittent dosing may not stably inhibit the NPM1c-XPO1 interaction, limiting its efficacy. In contrast, Eltanexor, currently tested in early-phase trials, can be administered more frequently (i.e. 5 days/week)^15^. Therefore, we asked whether prolonged XPO1 inhibition by Eltanexor could elicit a more pronounced anti-leukemic activity in *NPM1*-mutated cells.

## Methods

Selinexor and Eltanexor were evaluated in cellular and animal models of *NPM1*-mutated AML. Parental OCI-AML3, NPM1c-GFP OCI-AML3 (in-frame knock-in of GFP at the NPM1c endogenous locus)^7^ and NPM1c-FKBP(F36V)-GFP OCI-AML3 (in-frame knock-in of FKBP(F36V) and GFP at the NPM1c endogenous locus)^7^ cells were used for *in vitro* experiments. Patient-derived xenograft (PDX) of two *NPM1*-mutated AML patients were used for *in vivo* and *in vitro* experiments. RNA-sequencing data were analyzed applying the ARPIR pipeline and are available at GEO (GSE181176)^16^. Detailed methods are provided in the supplemental materials.

## Results and discussion

We first addressed the impact of intermittent XPO1 inhibition on the NPM1c-XPO1 interaction. As NPM1c subcellular localization is dependent on its binding to XPO1 (i.e. cytoplasmic when interacting with XPO1, nuclear when XPO1 is inhibited), we tracked the subcellular localization of endogenous NPM1c fused to GFP (NPM1c-GFP) in OCI-AML3 cells upon intermittent XPO1 inhibition. As expected, NPM1c-GFP was completely relocated to the nucleus after 12-hour Selinexor incubation. However, drug withdrawal caused cytoplasmic relocation over the following 24 hours, demonstrating quick recovery of the NPM1c-XPO1 interaction after transient XPO1 inhibition (Figure 1A).

**Figure 1.**
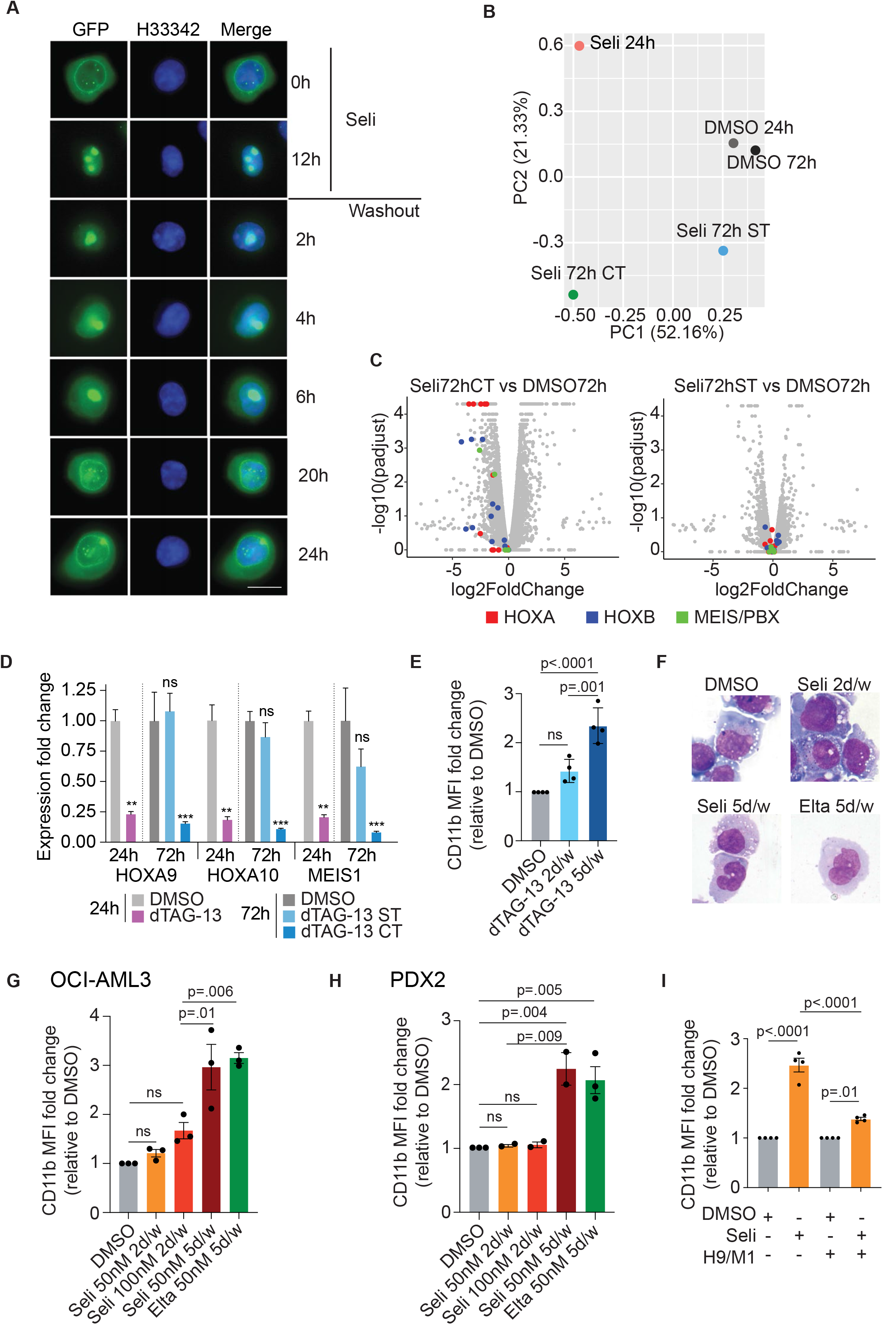
Prolonged XPO1 inhibition is necessary to elicit significant antileukemic activity in *NPM1*-mutated AML *in vitro.* **A)** Fluorescence microscopy of NPM1c-GFP OCI-AML3 treated for 12 hours with Selinexor 100nM. After 12 hours, Selinexor was removed from medium and cells left in culture for the following 24 hours taking pictures at 2, 4, 6, 20 and 24 hours after drug washout. Hoechst 33342 was used to stain nuclei of the cells. 100X magnification. Scale bar, 20 μm. **B)** Principal component analysis (PCA) plot derived from the means (N=2) of the FPKM values of parental OCI-AML3 cells collected at 24 hours treated with either DMSO or Selinexor 50 nM, and OCI-AML3 cells collected at 72 hours treated with either DMSO, Selinexor 50 nM short treatment (ST, 24h Selinexor + 48h fresh medium) or Selinexor 50 nM continuous treatment (CT, 72h Selinexor 50 nM). **C)** Volcano plots depicting differentially expressed genes in parental OCI-AML3 cells treated with 72h Selinexor continuous treatment (CT, 72h Selinexor 50 nM) and 72h Selinexor short treatment (ST, 24h Selinexor 50 nM + 48h fresh medium), compared to DMSO. Log_2_FC and Log_10_ p values are shown on the X and Y axis, respectively. Genes belonging to the HOXA (red) HOXB (blue) and MEIS/PBX (green) families are highlighted. **D)** *HOXA9, HOXA10 and MEIS1* expression by qPCR in NPM1c-FKBP(F36V)-GFP OCI-AML3 cells after 24 hours treatment with either DMSO or dTAG and 72h treatment with either DMSO, dTAG-13 ST (short treatment, 24h dTAG-13 + 48h fresh medium) or dTAG-13 CT (continuous treatment, 72h dTAG-13). N=3. Mean ± SEM. Tukey multiple comparison test. **E)** Flow-cytometry quantification of CD11b, expressed as MFI fold change relative to DMSO, in NPM1c-FKBP(F36V)-GFP OCI-AML3 cells at day 11 following treatment with either DMSO, dTAG-13 2 days/week or dTAG-13 5 days/week. N=4. Mean ± SEM. Tukey multiple comparison test. **F)** May-Grünwald Giemsa staining of OCI-AML3 cells at day 11 following treatment with either DMSO, Selinexor 2 days/week, Selinexor 5 days/week or Eltanexor 5 days/week. 40X magnification. **G)** Flow-cytometry quantification of CD11b, expressed as MFI fold change relative to DMSO, in OCI-AML3 cells at day 11 of treatment with either DMSO, Selinexor 50 nM 2 days/week, Selinexor 100 nM 2 days/week, Selinexor 50 nM 5 days/week and Eltanexor 50 nM 5 days/week. N=3. Mean ± SEM. Tukey multiple comparison test. **H)** Flow-cytometry quantification of CD11b, expressed as MFI fold change relative to DMSO, in PDX2 cells at day 11 of treatment with either DMSO, Selinexor 50 nM 2 days/week, Selinexor 100 nM 2 days/week, Selinexor 50 nM 5 days/week or Eltanexor 50 nM 5 days/week. N=3 for DMSO and Eltanexor, N=2 for the other groups. Mean ± SEM. Tukey multiple comparison test. **I)** Flow-cytometry quantification of CD11b, expressed as MFI fold change relative to DMSO, in untransduced and HOXA9/MEIS1-transduced OCI-AML3 cells after 7 days of continuous treatment with either DMSO or Selinexor 50nM. N=4. Mean ± SEM. Tukey multiple comparison test. H33342, Hoechst 33342; PC, principal component; FC, fold change; padjust, adjusted p value, MFI, Median Fluorescence Intensity; H9/M1, HOXA9/MEIS1; Seli, Selinexor. Elta, Eltanexor

As in *NPM1*-mutated cells high HOX/MEIS expression depends on the NPM1c-XPO1 interaction^7^, we hypothesized that early loss of XPO1 inhibition may result in inefficient HOX/MEIS downregulation. We determined *HOXA9, HOXA10* and *MEIS1* expression at 24 and 72 hours in cells treated with Selinexor for 24 hours (short treatment, ST) or for 72 hours (continuous treatment, CT). While ST caused only transient downregulation of HOX/MEIS expression, CT resulted in stable loss of these targets (supplemental Figure S1A). Next, to determine the impact of transient and stable XPO1 inhibition on the transcriptome, we performed RNA-sequencing in parental OCI-AML3 cells applying the same treatment strategy. After 24-hour incubation with Selinexor, the transcriptome was clearly perturbed, including downregulated HOX/MEIS (Figures 1B, supplemental Figures 1B, 1C, supplemental Table 1). However, drug withdrawal reduced transcriptional perturbation in the following 48 hours with only 57 residual differentially expressed genes (Figures 1B, 1C, supplemental Figures 1B, 1C, supplemental Table 1). Conversely, CT for 72 hours caused persistent downregulation of HOX/MEIS, combined with upregulation of genes related to myeloid differentiation and TP53 downstreams (Figures 1B, 1C, supplemental Figures 1C, 1D, supplemental Table 1). As XPO1 interacts with multiple cargo proteins^8^, to corroborate the hypothesis that the changes observed were mainly due to the loss of NPM1c-XPO1 interaction, we tested the effects of intermittent (2 days/week, e.g. Monday and Thursday) and prolonged (5 days/week, e.g. Monday to Friday) selective NPM1c degradation in CRISPR-engineered OCI-AML3 cells with endogenous NPM1c fused to the FKBP(F36V) degron tag and GFP^7^. Only prolonged NPM1c degradation caused stable HOX/MEIS downregulation at 72 hours and significant differentiation, mimicking what observed upon XPO1 inhibition (Figures 1D, 1E, supplemental Figures 1F, 1G). Altogether, these results clearly indicate that only prolonged loss of the NPM1c-XPO1 interaction can induce stable HOX/MEIS downregulation and differentiation in *NPM1*-mutated AML cells.

Next, we compared the ability of intermittent and prolonged XPO1 inhibition to induce differentiation of *NPM1*-mutated AML cells *in vitro.* Prolonged XPO1 inhibition with 50 nM of either Selinexor or Eltanexor induced differentiation of OCI-AML3 and *NPM1*-mutated PDX cells (PDX2)^7^, while 2 days/week treatment resulted in negligible changes (Figures 1F-1G, supplemental Figure S1H). Doubling the concentration of Selinexor to 100 nM 2 days/week did not significantly increase differentiation (Figure 1G). Importantly, ectopic expression of *HOXA9* and *MEIS1* significantly rescued differentiation upon prolonged XPO1 inhibition (Figure 1I, supplemental Figure S1I), confirming that persistent HOX/MEIS downregulation is required to achieve differentiation of *NPM1*-mutated cells.

Finally, we assessed the anti-leukemic potential of intermittent and prolonged XPO1 inhibition *in vivo* using two highly aggressive *NPM1/FLT3/DNMT3A* triple-mutated luciferase-expressing PDX models (PDX2 and PDX3). First, we treated NSG mice engrafted with PDX2 cells with either vehicle, Selinexor 2 days/week, Selinexor 5 days/week and Eltanexor 5 days/week (Figure 2A). HOX/MEIS expression and differentiation in sorted leukemic cells was analyzed after one week of treatment. While Selinexor 2 days/week did not induce changes of *HOXA9, HOXA10* and *MEIS1* nor of CD11b levels, both Selinexor and Eltanexor 5 days/week caused remarkable HOX/MEIS downregulation and differentiation (Figure 2B and 2C). Next, we investigated the leukemic engraftment of PDX2 cells by flow-cytometry and immunohistochemistry in the bone marrow following two weeks of treatment. Both 5 days/week regimens caused significant engraftment reduction, while 2 days/week Selinexor did not (Figure 2D and 2E), demonstrating that only prolonged XPO1 inhibition is effective against *NPM1*-mutated cells *in vivo.* Finally, to test the impact of prolonged XPO1 inhibition on AML growth *in vivo* and on survival, we treated both PDX2 and PDX3 mice with Eltanexor 5 days/week for 4 consecutive weeks. Eltanexor resulted in significant reduction of bioluminescence and prolonged survival compared to vehicle in both PDX models (Figures 2F-2I). Treatment was well tolerated with no progressive weight loss reported (supplemental Figures 2C and 2D).

**Figure 2:**
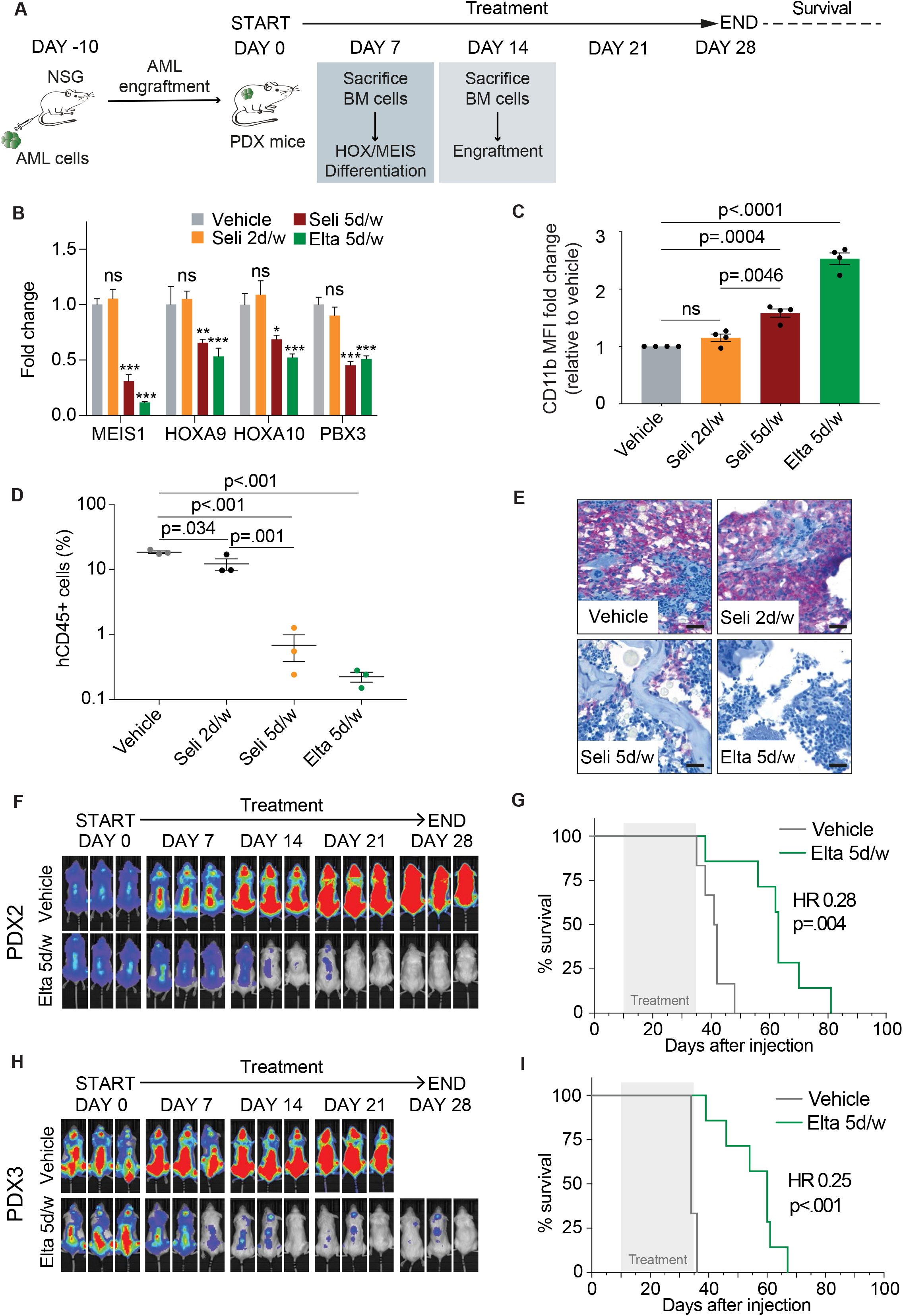
Prolonged XPO1 inhibition is necessary to elicit significant antileukemic activity in *NPM1*-mutated AML *in vivo.* **A)** Schematic overview of the *in vivo* experiments. Each NSG mouse was transplanted with 1×10^6^ GFP-Luc positive PDX cells. **B)** *HOXA9, HOXA10, MEIS1* and *PBX3* expression by qPCR in sorted PDX2 cells after 7 days of treatment with either vehicle, Selinexor 5 mg/kg 2 days/week, Selinexor 5 mg/kg 5 days/week and Eltanexor 10 mg/kg 5 days/week. N=4 mice per group. Mean ± SEM. Dunnett multiple comparison test. **C)** Flow-cytometry quantification of human CD11b, expressed as MFI fold change relative to vehicle, in sorted PDX2 cells after 7 days of treatment with either vehicle, Selinexor 5 mg/kg 2 days/week, Selinexor 5 mg/kg 5 days/week and Eltanexor 10 mg/kg 5 days/week. N=4 mice per group. Mean ± SEM. Tukey multiple comparison test. **D)** Bone marrow engraftment of PDX2 cells measured as human CD45 percent of positive cells after two weeks of treatment with either vehicle, Selinexor 5 mg/kg 2 days/week, Selinexor 5 mg/kg 5 days/week and Eltanexor 10 mg/kg 5 days/week. N=3 mice per group. Mean ± SEM. Tukey multiple comparison test. **E)** Representative images of BM histological sections stained for human CD45 after 14 days of treatment with either vehicle, Selinexor 5 mg/kg 2 days/week, Selinexor 5 mg/kg 5 days/week and Eltanexor 10 mg/kg 5 days/wee. 40X magnification. Scale bars, 20μm. **F)** Representative bioluminescence images of NSG mice transplanted with PDX2 cells treated with either vehicle (N=6) or Eltanexor 10 mg/kg (N=7) 5 days/week for 4 weeks. **G)** Kaplan-Meier curves of PDX2 mice treated with either vehicle (N=6) or Eltanexor 10 mg/kg (N=7) 5 days/week for 4 weeks. Treatment time is shown in light grey. Long-rank (Mantel-Cox) test. **H)** Representative bioluminescence images of NSG mice transplanted with PDX3 cells and treated with either vehicle (N=6) or Eltanexor 10 mg/kg (N=7) 5 days/week for 4 weeks. **I)** Kaplan-Meier curves of PDX3 mice treated with either vehicle (N=6) or Eltanexor 10 mg/kg (N=7) 5 days/week for 4 weeks. Treatment time is shown in light grey. Long-rank (Mantel-Cox) test. BM, bone marrow; Seli, Selinexor; Elta, Eltanexor; HR; hazard ratio.

*NPM1*-mutated AML is genetically well-characterized and is now included as distinct entity in the World Health Organization classification of myeloid neoplasms^17^. Therefore, there is growing interest in developing molecular targeted therapies for this AML variant^18^, including drugs interfering with HOX/MEIS expression, such as Menin-MLL^19–21^ and XPO1 inhibitors^7^. This study clearly demonstrates that 5 days/week XPO1 inhibition stably downregulates HOX/MEIS, induces differentiation and results into prolonged survival of *NPM1*-mutated PDX mice. In contrast, twice/week XPO1 inhibition does not elicit robust antileukemic activity *in vitro* and *in vivo* in *NPM1*-mutated AML, likely explaining the lack of benefit of Selinexor in patients with this leukemia. How XPO1 inhibition results in HOX/MEIS downregulation remains unclear. Possible hypotheses include nuclear relocation of NPM1c interactors with transcriptional repressive properties^2^ and displacement of NPM1c from XPO1 bound at HOX/MEIS loci^2,22^. Furthermore, mechanisms other than those mediated by NPM1c, e.g. TP53 activation (supplemental Figure 1D and ^9^), may contribute to the anti-leukemic effects of XPO1 inhibitors in *NPM1*-mutated cells. In conclusion, as Phase 1 data of Eltanexor have shown it can be safely administered 5 days/week^15^, this study lays the groundwork for the appropriate design of new clinical trials with XPO1 inhibitors in *NPM1*-mutated AML.

## Acknowledgments

This work has been supported by the Associazione Italiana per la Ricerca sul Cancro (AIRC Start-Up Grant 2019 n. 22895) and the European Research Council (ERC Adv grant 2016 n. 740230 and ERC Cons Grant 2016 n. 725725).

## Authorship contribution

G.P., B.F. and L.B. conceived the study. G.P., A.G., F.R. and A.M. performed *in vitro* experiments. B.B. performed IHC analysis. G.P. and F.M. performed *in vivo* experiments. S.C. and C.Q. performed RNA-sequencing. V.T. and G.S. performed bioinformatic analysis. M.P.M., P.S., F.L., S.S. and Y.L. provided reagents and critical inputs. G.P., B.F. and L.B. analyzed the results and wrote the manuscript with the input from all the authors.

