## Supplemental materials for "Prolonged XPO1 inhibition is essential for optimal anti-leukemic activity in *NPM1*-mutated AML"

### Supplemental methods

*AML cell lines*

Parental OCI-AML3 cells (ATCC), harboring NPM1 mutation A, GFP-tagged OCI-AML3 cells (NPM1c-GFP OCI-AML3), bearing NPM1 mutant allele fused to GFP and FKBP-based degron-tagged OCI-AML3 cells (NPM1c-FKBP(F36V)-GFP OCI-AML3), holding NPM1 mutant allele fused to FKBP12 (F36V) and GFP, were cultured in RPMI-1640 medium supplemented with 10% Fetal Bovine Serum (FBS), 1% L-glutamine (Glu) and 1% penicillin/streptomycin (P/S). All cell lines were grown in mycoplasma-free conditions and maintained in 5% $CO_2$ at 37°C at a concentration of 0.5-1.5x$10^6$ cells/ml.

*Patient-derived xenografts (PDXs)*

Leukemic cells from two *NPM1/FLT3/DNMT3A* triple mutated AML patients (PDX2 and PDX3) were injected subcutaneously into NSG mice, leading to the formation of solid masses at the site of injection. Mice were sacrificed and tumors were isolated, crushed and filtered to obtain a single cell suspension. An aliquot of cells was used for the experiments and leftover cells were subcutaneously injected into new NSG recipient mice, allowing for propagation. This protocol was approved by the Ethics Committee of the University of Perugia and patients gave written consent for the use of the cells. To obtain PDX2 cells for *in vitro* experiments, subcutaneous tumors were isolated, crushed and filtered to obtain a single cell suspension. Viable cells were resuspended in IMDM medium supplemented with 10% FBS, 1% Glu, 1% P/S and cytokines (30ng/ml FLT3L, 30ng/ml G-SCF and 15ng/ml TPO (all Cell Guidance Systems) and maintained at a concentration of 2.0x$10^6$ cells/ml.

*Treatment of OCI-AML3 NPM1c-GFP cells with Selinexor and live cell imaging*

NPM1c-GFP OCI-AML3 cells were treated with 100nM Selinexor for 12 hours. Cells were harvested, washed twice in PBS, resuspended in fresh medium (referred as drug washout) and left in culture for the following 24 hours while taking live pictures. Live cell fluorescence images were collected with the Olympus IX51 microscope equipped with a 100X/1.30 oil immersion objective and processed with the CellSens Digital Imaging Software (Olympus). Vital dye Hoechst 33342 (Thermo Fisher Scientific) was used to stain nuclei of the cells.

*Treatment of OCI-AML3 NPM1c-GFP and parental OCI-AML3 cells with Selinexor*

OCI-AML3 NPM1c-GFP and parental OCI-AML3 cells were treated with 50nM Selinexor or 0.3% DMSO (control) for 24 hours (short treatment – ST) followed by drug washout from the medium or continuously for 72 hours (continuous treatment – CT). After 24 and 72 hours of treatments cells were harvested for total RNA purification and used for further gene expression analyses by qRT-PCR or RNA-sequencing studies.

*Treatment of* NPM1c-FKBP(F36V)-GFP OCI-AML3 *with dTAG-13*

To study the impact of intermittent (2 days/week, i.e. Monday and Thursday) and prolonged (5 days/week, i.e. Monday to Friday) selective NPM1c degradation NPM1c-FKBP(F36V)-GFP OCI-AML3 cells were plated at $0.5x10^6$ cells/ml and treated with 500nM dTAG-13 (Tocris Bioscience) or 0.3% DMSO as a control and cultured for 11 days. Cells were replated at the same concentration in fresh medium with or without dTAG-13 every 24 to 72 hours, as needed. Differentiation was assessed at day 11 by flow cytometry. $0.5x10^6$ cells were collected at 72 hours in RNA lysis buffer for total RNA purification and subsequent analyses.

*Intermittent and prolonged XPO1 inhibition in parental OCI-AML3 and PDX2 cells*

To study the effect of intermittent (2 days/week, i.e. Monday and Thursday) and prolonged (5 days/week, , i.e. Monday to Friday) XPO1 inhibition on differentiation of OCI-AML3 and PDX2, cells were plated at $0.5x10^6$ or $2.0x10^6$ cells/ml, respectively and treated with 50nM Selinexor 2 days/week, 100nM Selinexor 2 days/week, 50 nM Selinexor 5 days/week, 50 nM Eltanexor 5 days/week or 0.3% DMSO as a control and cultured for 11 days. Cells were replated at the same concentration in fresh medium with or without XPO1 inhibitors every 24 to 72 hours, as needed. Differentiation was assessed by flow cytometry at day 11.

*Lentivirus production and lentiviral transduction*

The lentiviral vector pLV[Exp]-Puro-EFS>hHOXA9[NM_152739.3](ns):T2A:hMEIS1 [NM_002398.2] used to ectopically express *HOXA9* and *MEIS1* in OCI-AML3 cells was purchased from VectorBuilder (vector ID: VB170808-1055qbq). Virus particles were produced by transient transfection of sub-confluent 293FT cells (ThermoFisher Scientific) with the packaging plasmid psPAX2 (Addgene plasmid #12260) and the envelope plasmid pMD2.G (Addgene plasmid #12259) using Lipofectamine 2000 (ThermoFisher Scientific). Viral supernatant was collected 48 hours and 72 hours after transfection and filtered (0.45 mm). OCI-AML3 cells were infected by direct addiction of viral supernatant and polybrene (8 ug/ml) onto 6-well cell culture plates. Selection of transduced cells started 72h after infection by addition of 1 mg/ml puromycin (Sigma-Aldrich) to the medium. Parental untransduced OCI-AML3 cells were used as control for puromycin selection efficiency. High *HOXA9* and *MEIS1* mRNA levels were confirmed by qRT-PCR analysis.

*Flow cytometry*

In each experiment, $2x10^5$ cells were washed with PBS and stained for 20 min at 4°C in the dark with the following antibodies: PE-conjugated anti-CD11b (BD Bioscence) were used to assess differentiation levels. FITC-conjugated anti-CD45 (BD Bioscence) was used to assess AML engraftment. Excess antibody was removed by washing the cells with PBS. In all samples, cell debris were excluded based on FSC-A/SSC-A dot-plot and 7-Amino-Actinomycin D (7-AAD, BD-Bioscience) was used for dead-cell exclusion. Singlets were gated using the FSC-A/FSC-H dot-plot. Flow cytometry and cell sorting were performed respectively with the BD FACSCanto II and the BD FACSAria III Sorter. Flow cytometry data were analyzed with FlowJo 10.7 software (BD Bioscence) and expressed as fold change of the median fluorescence intensity (MFI) of the treated sample, relative to the control one.

*In vivo studies*

All animal procedures were performed in accordance with the European Community guidelines, and approved by the Institutional Animal Care Committee of University of Perugia and the Italian Ministry of Health (Authorization n. 104/2020-PR). 8- to 10-week-old female NOD.Cg-Prkdc$^{scid}$Il2rg$^{tm1Wjl}$/SzJ (NSG) mice (Charles River Europe) were intravenously injected with $1x10^6$ primary AML cells (PDX2 and PDX3) stably transduced with the pHIV-Luc-ZsGreen vector (Addgene, plasmid #39196) that co-expresses firefly luciferase (Luc2P) and ZsGreen protein under the human EF-1alpha promoter. For one-week and two-weeks end point experiments with PDX2 model, at AML engraftment 10 days post-transplantation (luminescent flux $>10^6$ as assessed by basal bioluminescence (BLI)) mice were split into vehicle (N=4 and N=3), Selinexor 2 days/week (Seli 2 d/w, e.g. Monday and Thursday, N=4 and N=3), Selinexor 5 days/week (Seli 5 d/w, e.g. Monday to Friday, N=4 and N=3) and Eltanexor 5 days/week (Elta 5d/w, e.g. Monday to Friday, N=4 and N=3) treatment groups and dosed by oral gavage (p.o.) with vehicle (1% Tween-80/0.5% methyl cellulose), Selinexor (5 mg/kg) or Eltanexor (10 mg/kg) for one or two weeks, respectively. Mice were sacrificed at day 7 or 14 and bone marrow (BM) cells were isolated by crushing both tibias and femurs in 1X PBS supplemented with 5% FBS. Leukemic engraftment was monitored by flow cytometry analysis of human CD45. ZsGreen-positive cells were sorted using the BD FACS Aria III cell sorter and used for qRT-PCR analysis of HOX/MEIS expression and flow cytometry analysis of human CD11b to evaluate cell differentiation. Sorting purity was ≥85%. In addition, spines were collected for immunohistochemical analysis of human CD45. For survival experiments with PDX2 and PDX3 models, after AML engraftment, mice were split into vehicle (N=6) and Elta 5d/w (N=7) treatment groups and dosed with vehicle (1% Tween-80/0.5% methyl cellulose; p.o.) or Eltanexor (10 mg/kg; p.o.) 5 times a week (e.g. Monday to Friday) for four weeks. Body weight was monitored twice weekly and leukemic progression was monitored weekly by BLI measurement: mice were subcutaneously injected with luciferin (XenoLight D-Luciferin Potassium Salt, PerkinElmer) and kept anesthetized with isoflurane before *in vivo* imaging with the IVIS Lumina III In Vivo Imaging System. Quantification and image processing were performed using the Living Imaging Software (PerkinElmer).

*Bone marrow samples and immunohistochemistry (IHC) analysis*

Spines were fixed in formalin, decalcified in Osteodec (Bio-optica), dehydrated and then processed for paraffin embedding. Paraffin-embedded tissues were cut into 3-mm-thick sections and subjected to antigen retrieval. Unmasking was performed by incubating the paraffin sections in EnVision FLEX Target Retrieval Solution High pH (Dako-Agilent) at 85°C for 5 minutes. Sections were then immunostained for 30 min at room temperature with human anti-CD45 antibody (Monoclonal mouse Anti-Human CD45, Leucocyte Common Antigen Clones 2B11 + PD7/26; Dako) diluted in Dako REAL Antibody diluent (Dako-Agilent) and processed in a Dako Autostainer using the Dako REAL Detection System, Alkaline Phosphatase/RED, Rabbit/Mouse (Dako-Agilent). Sections were then counterstained in hematoxylin, Mayer's (Dako-Agilent) for 5 minutes and mounted in Kaiser's glycerol gelatin (Merck Millipore). Immunohistochemical stain pictures were acquired with a U Plan FLN 40X/0.75

objective of an Olympus BX51 microscope equipped with an Olympus DP71 digital camera, using the TC Capture acquisition software.

*RNA isolation and qRT-PCR gene expression analysis*

Total RNA was isolated according to manufacturer's instructions using the Quick-RNA Microprep Kit (Zymo Research) including in-column DNase I treatment. RNA quality control and quantification was performed using the NanoDrop spectrophotometer. RNA was reverse transcribed with the SuperScript IV First-Strand Synthesis System Kit (Invitrogen). RT-PCR was performed on a CFX96 Real-Time PCR system (Bio-Rad) using the iQ SYBR Green Supermix (Bio-Rad) and custom primers. *GAPDH* was used as the housekeeping gene for normalization. Relative gene expression and fold change were calculated by the $2^{-\Delta\Delta Ct}$ method. All the primers are listed as follow: *HOXA9* (FW: 5'-GACAAGCCCCCCATCGAT-3', REV: 5'-GTGGAGCGCGCATGAAG-3'); *HOXA10* (FW: 5'-AAAGCCTCGCCGGAGAA-3', REV: 5'-GTTGGCTGCGTTTTCACCTT-3'); *MEIS1* (FW: 5'-CGTGGCTGTTCCAGCATCTA-3', REV: 5'-GCCAACTGCTTTTTCTGTTCTTC-3') and *GAPDH* (FW: 5'-TTTTGCGTCGCCAGCCGAG-3', REV: 5'-GGCGCCCAATACGACCAAAT-3').

*RNA sequencing and data analysis*

RNA was extracted from two replicates for each condition. RNA quality and quantity was assesses using a Nanodrop spectrophotometer. Preparation of sequencing libraries was conducted with 250 ng of total RNA from each sample using the TruSeq Stranded mRNA Library Prep Kit (Illumina) following standard procedures. Amplified libraries were purified and quantified using the KAPA Library Quantification Kit (Roche). Single-end sequencing (75bp) was conducted on an Illumina NextSeq500 instrument. After RNA sequencing, FastQ files were generated, demultiplexed and the lanes produced for each sample were combined into a single file. Analysis and comparison of the samples was performed with the ARPIR pipeline [1] that uses HISAT2 for alignment and Cufflinks for quantification and differential analysis. Initial quality control on FastQ files using FastQC and FastQ Screen was followed by a pre-processing phase aimed at eliminating the reads belonging to contaminating RNA and in particular PhiX genome and ribosomal RNA. The FastQs obtained were aligned with HISAT2 to obtain the BAM files, which underwent a second quality control via RSeQC.

After the quality check and the alignment (to hg19), Samtools was used to sort and index the BAM files. Cufflinks was then used to generate transcript abundance and differential expression analysis (DEA) files. To eliminate the zeros and make the fold change readable in logarithmic format, 0.25 has been added to all the values of fragments per kilobase million (FPKM). After the classical DEA, gene set enrichment analysis [2,3] (4.1 version) was performed on the differentially expressed genes (with $log_2$Fold Change value ≥1.5 and adjusted p-value <0.05). The principal component analysis (PCA) plot, obtained with the autoplot and prcomp functions in R, was created starting from the FPKM data, while the volcano plot was obtained with the R package ggplot2. The heatmap in supplemental Figure 1 was generated using Morpheus (https://software.broadinstitute.org/morpheus/).

*Statistical analysis*

Graphs generation and statistical analyses were performed using Prism 8 (GraphPad). Student's t-test or One-way ANOVA or 2-way ANOVA test and their relative post-hoc multicomparison tests (i.e. Dunnett and Tukey) were used to calculate the significance level between groups, as indicated in figure legends. Log-rank (Mantel-Cox) test was used to assess significance in survival experiments. P values of less than 0.05 were considered significant. The number of independent biological replicates is reported in each figure legend.

*Data and software availability*

Raw and processed RNA-seq data have been deposited to the NCBI Gene Expression Omnibus (GEO: GSE181176). Software programs used in this study were from publicly available resources.

[2] Subramanian A, Tamayo P, Mootha VK, Mukherjee S, Ebert BL, Gillette MA, Paulovich A, Pomeroy SL, Golub TR, Lander ES, Mesirov JP. Gene set enrichment

analysis: a knowledge-based approach for interpreting genome-wide expression profiles. Proc Natl Acad Sci U S A. 2005 Oct 25;102(43):15545-50.

[3] Mootha VK, Lindgren CM, Eriksson KF, Subramanian A, Sihag S, Lehar J, Puigserver P, Carlsson E, Ridderstråle M, Laurila E, Houstis N, Daly MJ, Patterson N, Mesirov JP, Golub TR, Tamayo P, Spiegelman B, Lander ES, Hirschhorn JN, Altshuler D, Groop LC. PGC-1alpha-responsive genes involved in oxidative phosphorylation are coordinately downregulated in human diabetes. Nat Genet. 2003 Jul;34(3):267-73.

**Supplemental Table 1. RNA-sequencing data.** FPKMs, log2Fold Change and p values for all samples analyzed.

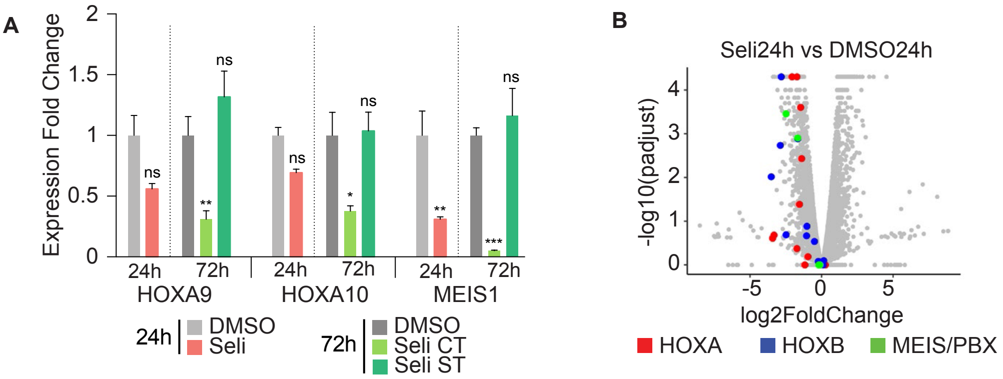


C

| | Genes Up* | Genes Down^ |
|---|---|---|
| 24 hours | 131 | 279 |
| 72 hours CT | 225 | 263 |
| 72 hours ST | 21 | 26 |

*$log_2$FC vs DMSO ≥ 1.5; p value ≤ 0.05
^$log_2$FC vs DMSO ≤ -1.5: p value ≤ 0.05

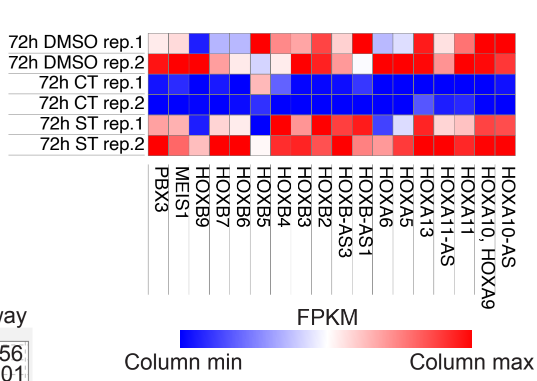


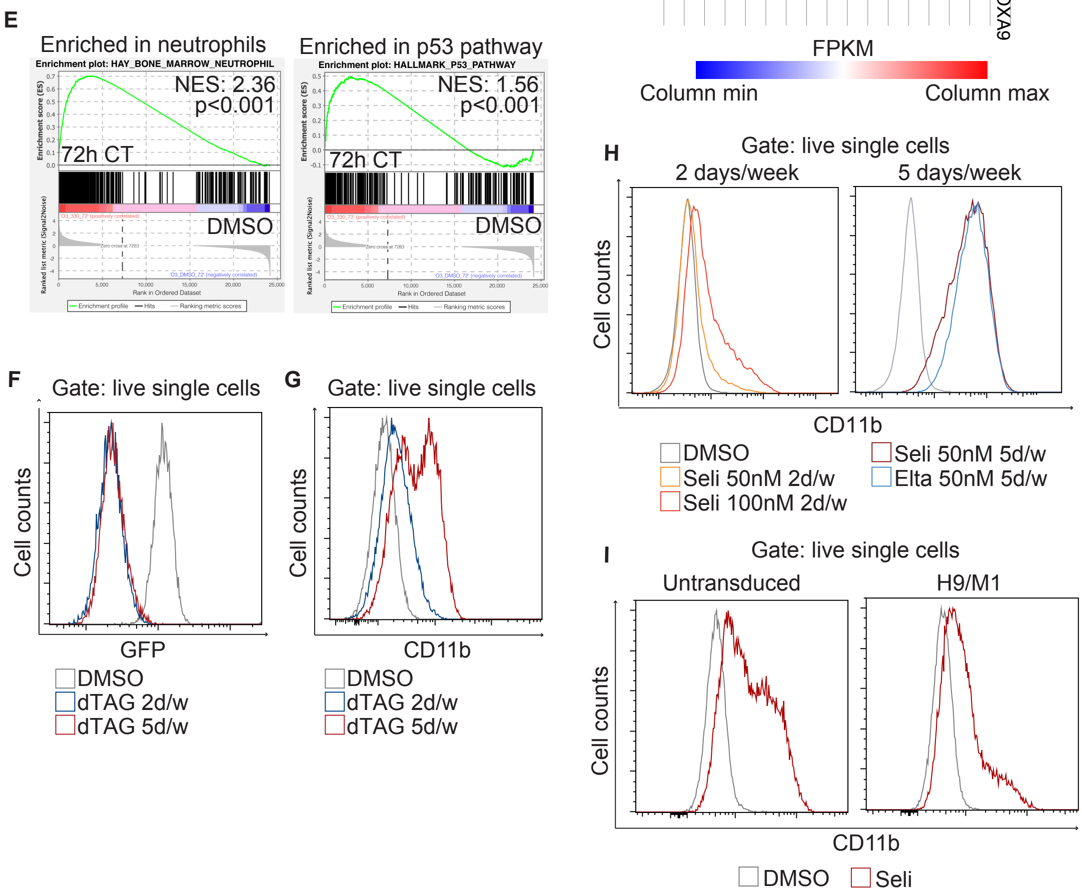


Supplemental Figure 1

**Supplemental Figure 1. A)** *HOXA9, HOXA10 and MEIS1* expression by qPCR in NPM1c-GFP OCI-AML3 cells after 24 hours treatment with either DMSO or Selinexor and 72h treatment with either DMSO, Selinexor ST (short treatment, 24h Selinexor + 48h fresh medium) or Selinexor CT (continuous treatment, 72h Selinexor). N=3. Mean $\pm$ SEM. Tukey multiple comparison test. **B)** Volcano plot depicting differentially expressed genes in OCI-AML3 cells treated with 24h Selinexor versus DMSO. $Log_2$FC and $Log_{10}$p-value are shown on the X and Y axis, respectively. Genes belonging to the HOXA (red) HOXB (blue) and MEIS/PBX (green) families are highlighted. N=2. **C)** Upregulated ($log_2$FC $\geq$ 1.5 and p value $\leq$ 0.05) and downregulated ($log_2$FC $\leq$ 1.5 and p value $\leq$ 0.05) genes in RNA-sequencing data from parental OCI-AML3 cells treated for 24 hours with either DMSO or 50 nM Selinexor, or 72 hours with either DMSO, Selinexor ST (short treatment, 24h Selinexor + 48h fresh medium) or Selinexor CT (continuous treatment, 72h Selinexor). N=2. **D)** Heatmap from RNA-sequencing data depicting expression levels of *HOXA, HOXB, MEIS1* and *PBX3* genes in parental OCI-AML3 cells treated for 72h with either DMSO, Selinexor CT (continuous treatment, 72h Selinexor) or Selinexor ST (short treatment, 24h Selinexor + 48h fresh medium). N=2. **E)** Gene set enrichment analysis from RNA-sequencing data of parental OCI-AML3 cells treated for 72 hours with Selinexor CT or DMSO. Enrichment plots for "Hay_bone_marrow_neutrophils" and "Hallmark_P53_pathway" gene sets are shown. **F)** Flow cytometry analysis of NPM1c-FKBP(F36V)-GFP OCI-AML3 cells treated with dTAG-13. NPM1c-GFP levels demonstrate efficient degradation of NPM1c upon dTAG-13. **G)** Histogram plot representing CD11b expression analyzed by flow cytometry at day 11 in NPM1c-FKBP(F36V)-GFP OCI-AML3 cells treated with either DMSO, dTAG-13 2 days/week or dTAG-13 5 days/week. **H)** Histogram plot representing CD11b expression analyzed by flow cytometry at day 11 in OCI-AML3 cells treated with either DMSO, Selinexor 50 nM 2 days/week, Selinexor 100 nM 2 days/week, Selinexor 50 nM 5 days/week or Eltanexor 50 nM 5 days/week. **I)** Histogram plot showing CD11b expression analyzed by flow cytometry at day 7 in OCI-AML3 cells or OCI-AML3 cells transduced with *HOXA9/MEIS1* lentivirus and treated with either DMSO or Selinexor 50 nM for 7 consecutive days.

Seli, Selinexor; Elta, Eltanexor; FC, fold change; padjust, adjusted p value; rep., replicate; NES, Normalized Enrichment Score.

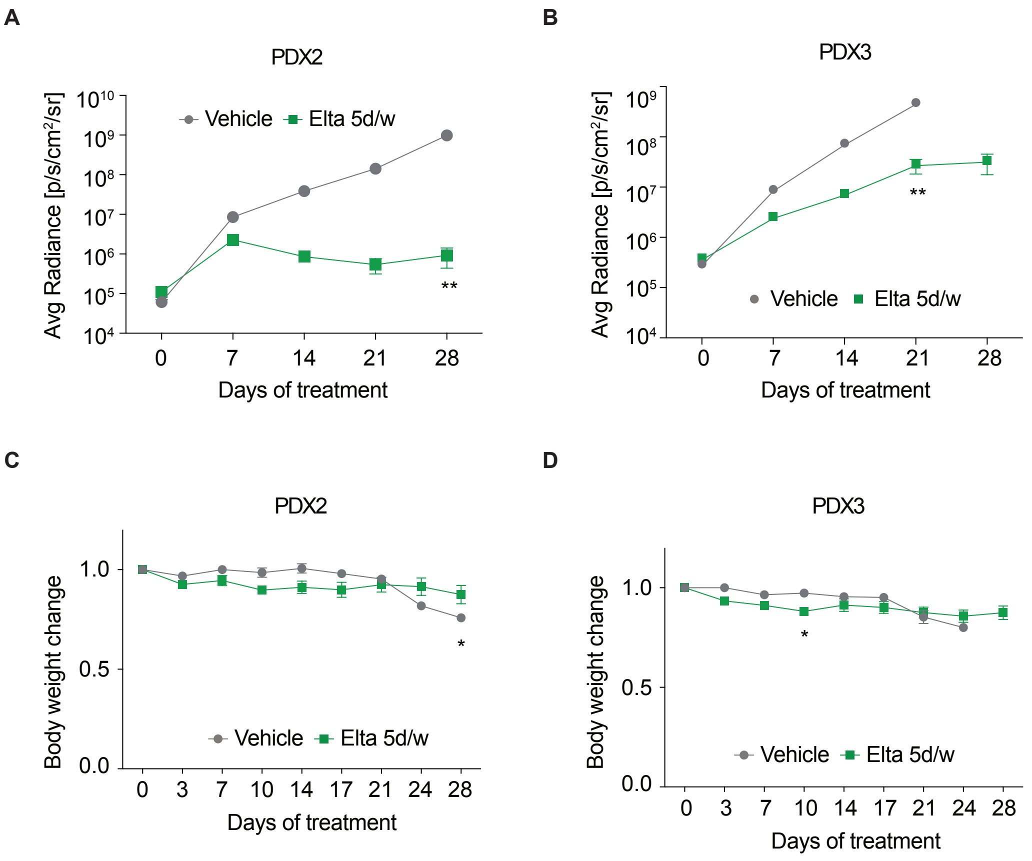


Supplemental Figure 2

**Supplemental Figure 2. A-B)** Bioluminescence level, expressed as average radiance (p/s/cm$^2$/sr), over time for each animal shown in figure 2F (PDX2) and 2H (PDX3), treated with either vehicle of Eltanexor 10 mg/kg 5 days/week. Mean ± SEM, Mann-Whitney test. **C-D)** Body weight changes of PDX2 and PDX3 mice over 28 days of treatment with either vehicle of Eltanexor 10 mg/kg 5 days/week. Body weight measurements were taken twice weekly and are referenced to the values obtained at the first day of treatment, which was set to 1. Mean ± SEM, Sidak multiple comparison test. Elta, Eltanexor.
